# Predicting functional effect of missense variants using graph attention neural networks

**DOI:** 10.1101/2021.04.22.441037

**Authors:** Haicang Zhang, Michelle S. Xu, Wendy K. Chung, Yufeng Shen

## Abstract

Accurate prediction of damaging missense variants is critically important for interpreting genome sequence. While many methods have been developed, their performance has been limited. Recent progress in machine learning and availability of large-scale population genomic sequencing data provide new opportunities to significantly improve computational predictions. Here we describe gMVP, a new method based on graph attention neural networks. Its main component is a graph with nodes capturing predictive features of amino acids and edges weighted by coevolution strength, which enables effective pooling of information from local protein context and functionally correlated distal positions. Evaluated by deep mutational scan data, gMVP outperforms published methods in identifying damaging variants in *TP53, PTEN, BRCA1*, and *MSH2*. Additionally, it achieves the best separation of *de novo* missense variants in neurodevelopmental disorder cases from the ones in controls. Finally, the model supports transfer learning to optimize gain- and loss-of-function predictions in sodium and calcium channels. In summary, we demonstrate that gMVP can improve interpretation of missense variants in clinical testing and genetic studies.

## Main

Missense variants are major contributors to genetic risk of cancers ^1,2^ and developmental disorders ^3-5^. Missense variants have been used, along with protein-truncating variants, to implicate new risk genes and are responsible for many clinical genetic diagnoses. However, the majority of rare missense variants are likely benign or only have minimal functional impact. As a result of the uncertainty of the functional impact, most rare missense variants reported in clinical genetic testing are classified as variants of uncertain significance (VUS)^6^, leading to ambiguity, confusion, overtreatment, and missed opportunities for clinical intervention. In human genetic studies to identify new risk genes by rare variants, pre-selecting damaging missense variants based on computational prediction is a necessary step to improve statistical power ^4,5,7,8^. Therefore, computational methods are critically important to interpret missense variants in clinical genetics and disease gene discovery studies.

Numerous methods, such as Polyphen ^9^, SIFT ^10^, CADD^11^, REVEL^12^, MetaSVM^13^, M-CAP^14^, Eigen^15^, MVP^16^, PrimateAI^17^, MPC^18^, and CCRs^19^, have been developed to address the problem. These methods differ in several aspects, including the prediction features, how the features are represented in the model, the training data sets, and how the model is trained. Sequence conservation or local protein structural properties are the main prediction features for early computational methods such as GERP^20^ and PolyPhen. MPC and CCRs estimate sub-genic coding constraints from large human population sequencing data which provide additional information not captured by previous methods. PrimateAI learns protein context from sequences and local structural properties using deep representation learning. A number of studies have reported evidence that functionally damaging missense variants are clustered in 3-dimensional protein structures^21-23^.

Here we present gMVP, a graph attention neural network model designed to effectively represent or learn the representation of all the information sources to improve prediction of functional impact of missense variants. gMVP uses a graph to represent a variant and its protein context with node features describing sequence conservation and local structural properties. gMVP uses a graph attention neural network to learn the representation of a large protein context, and uses coevolution strength as edge features which can potentially pool information about conservation and coding constraints of distal but functionally correlated positions. We trained gMVP using curated pathogenic variants and random rare missense variants in human population. We then benchmarked the performance using data sets that have been curated or collected by entirely different approaches, including cancer somatic mutation hotspots ^24^, functional readout datasets from deep mutational scan studies of well-known risk genes^25-28^, and *de novo* missense variants from studies of autism spectrum disorder (ASD) ^4^ and neurodevelopmental disorder (NDD)^5^. Finally, we investigated the potential utility of transfer learning for classifying gain- and loss-of-function variants in specific gene families based on the generic model trained across all genes.

## Results

### Model architecture and prediction features

gMVP is a supervised machine learning method for predicting functionally damaging missense variants. The functional consequence of missense variants depends on both the type of amino acid substitution and its protein context. gMVP uses a graph attention neural network to learn representation of protein sequence and structure context and context-dependent impact of amino acid substitutions on protein function.

The main component of gMVP is a graph that represents a variant and its protein context (Figure 1 and Supplementary Figure 1). Given a variant, we define the 128 amino acids flanking the reference amino acid as protein context. We build a star-like graph with the reference amino acid as the center node and the flanking amino acids as context nodes and connect the center node and every context node with edges. We use coevolution strength between the center node of variant and the context node as edge features. Coevolution strength is highly correlated with functional interactions and protein residue-residue contact that captures the potential 3D neighbors in folded proteins^29-32^. For the center node, we include as features the amino acid substitution, evolutionary sequence conservation, and predicted local structural properties such as secondary structures (Methods). For context nodes, in addition to primary sequence, sequence conservation, and local structure features, we also include expected and observed number of rare missense variants in human population to capture selection effect of damaging variants in human^18,19^. Let ***x***, {***n***_*i*_}, and {***f***_*i*_} denote input feature vectors for the center node, neighbor nodes, and edges, respectively. We first use three 1-depth dense layers to encode ***x***, {***n***_*i*_}, and {***f***_*i*_} to latent representation vectors ***h***, {***t***_***i***_}, and {***e***_***i***_}, respectively. We then use a multi-head attention layer to learn attention weight ***w***_***i***_ for each neighbor and to learn a context vector ***c*** by weighting the neighbors. Attention scores play a key part in attention-based neural networks^33,34^. Our attention scores account for both the node features and the edge features. Specifically, we use *tanh* (***W***[***h, t*_*i*_**, ***e*_*i*_**]) as attention scores where *tanh* denotes a hyperbolic tangent activation function, where ***W*** is the weight matrix to be trained. Next, we used a recurrent neural layer^35^, which is widely used to leverage sequence context in natural language modelling, to integrate the context vector ***c*** and the vector ***h*** of variant. Finally, we use a linear projection layer and a sigmoid layer to perform classification and output the damaging scores.

**Figure 1.**
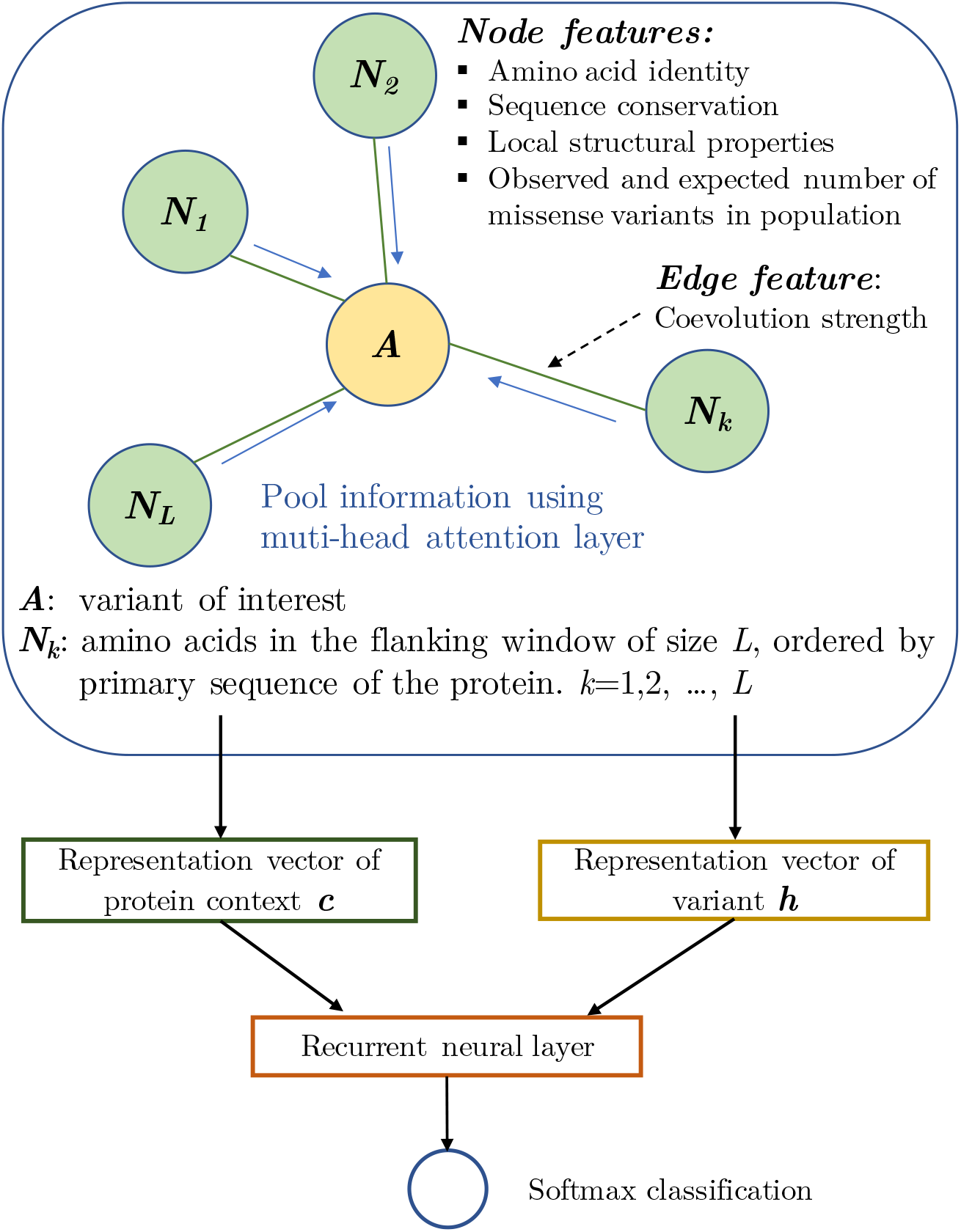
An overview of gMVP model. gMVP uses a graph to represent a variant and its protein context defined as 128 amino acids flanking the reference amino acid. The amino acid of interest is the center node (colored as orange) and the flanking amino acids are the context nodes (colored as light green). All context nodes are connected with the center node but not each other. The edge feature is coevolution strength. The node features include conservation and predicted structural properties. Additionally, center node features include the amino acid substitution; context node features include the primary sequence and the expected and observed number of rare missense variants in human population. We use three 1-depth dense layers to encode the input features to latent representation vectors and used a multi-head attention layer to learn a context vector ***c***. We then use a recurrent neural layer connected with softmax layer to generate prediction score from the context vector ***c*** and the representation vector ***h*** of variant.

### Model training and testing

We collected likely pathogenic and benign missense variants from curated databases (HGMD^36^, ClinVar^37^, and UniProt^38^) as training positives and negatives, respectively, excluding the variants with conflicting evidence in the databases (see Methods). To balance positive and negative sets, we randomly selected rare missense variants observed in human population sequencing data DiscovEHR as additional negatives for training. In total there are 59,701 positives and 59,701 negatives, which cover 3,463 and 14,222 genes, respectively. We used stochastic gradient descent algorithm^39^ to update the model’s parameters with an initial learning rate of 1e-3, and applied early stopping with validation loss as metric to avoid overfitting. We implemented the model and training algorithms using TensorFlow^40^. Running on a Linux workstation with 1 NVIDIA Titan RTX GPU, the whole training process took ∼4 hours. When benchmarking the performance using a range of datasets, we compared gMVP with other widely used methods in genetic studies including PrimateAI^17^, M-CAP^41^, CADD^11^, MPC^18^, and REVEL^12^.

Human-curated pathogenic variants have hidden false positives that are likely caused by systematic bias and errors, which can be picked up by deep neural networks. Therefore, conventional approaches for performance evaluation using testing data randomly partitioned from the same source as training data usually lead to inflated performance measure. To objectively evaluate the performance of the model, we compiled cancer somatic mutations that are unlikely to share the same systematic errors as the training data sets. We included missense mutations located in inferred hotspots based on statistical evidence from a recent study ^24^ as positives and randomly selected rare variants from DiscovEHR database^42^ as negatives. The gMVP score distributions of cancer hotspot mutations and random variants have distinct modes (Figure 2a). When compared to published methods, gMVP achieved the best performance with an area under the receiver operating characteristic curve (AUROC) of 0.88 (Figure 2b and Supplementary Table 2). REVEL is close with an AUROC of 0.86.

**Figure 2.**
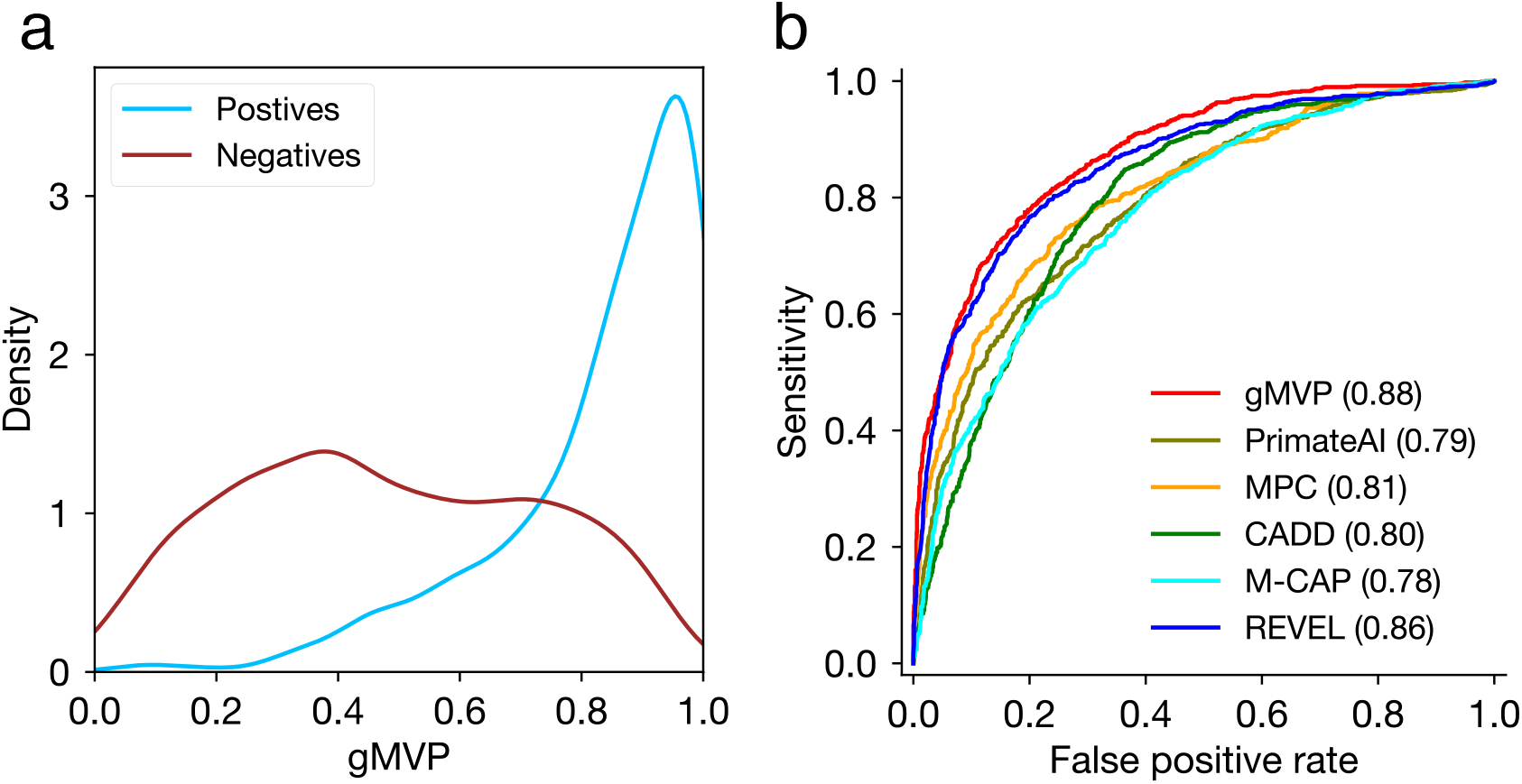
Evaluating gMVP and published methods using cancer somatic mutation hotspots and random variants in population. (a) The gMVP score distributions for variants in cancer hotspots (labeled positives) and random missense variants in population (labeled negatives). (b) Comparison of ROC curves of gMVP and published methods. The ROC curves are evaluated on 878 cancer mutations located in hotspots from 209 genes, and 1756 (2 times of the positives) randomly selected rare variants from the DiscovEHR data.

### gMVP can identify damaging variants in known disease genes

Missense variants that occur in different protein contexts, even in the same gene, can have different consequences. This is the core problem of interpretation of variants from known risk genes in clinical genetic testing and discovery of new disease genes. As performance evaluation using variants across genes are confounded by gene-level properties, here we aim to evaluate gMVP and other methods in distinguishing damaging variants from neutral variants in the same genes. To this end, we obtained functional readout data from deep mutational scan assays of four well-known disease risk genes, *TP53*^*28*^, *PTEN*^*27*^, *BRCA1*^26^, and *MSH2*^25^, as benchmark data. The data includes 432 damaging (“positives”) and neutral (“negatives”) 1,476 negatives for *BRCA1*, 262 positives and 1632 negatives for *PTEN*, 540 positives and 1,108 negatives for *TP53*, and 414 positives and 5439 negatives for *MSH2*, respectively. We note that during gMVP training, all variants in these four genes were excluded to avoid inflation in performance evaluation.

We first investigated the gMVP score distributions of damaging and neutral variants. Damaging variants clearly have different score distribution compared to the neutral ones in each gene (Supplementary Fig. 2). Additionally, gMVP scores are highly correlated with functional scores from the deep mutational scan assays, with a Spearman correlation coefficient of 0.67 (*p*=1e-246), −0.48 (*p*=8e-122), −0.53 (*p*=7e-151), and 0.29 (*p*=7e-117) in *TP53, PTEN, BRCA1* and *MSH2*, respectively (Supplementary Fig. 3 and Supplementary Table 3-6).

We then used functional readout data as ground truth to estimate precision/recall and compared gMVP with other methods. The areas under the precision-recall curves (AUPRC) of gMVP are 0.78, 0.85, 0.81, and 0.39 for *PTEN, TP53, BRCA1*, and *MSH2*, respectively (Figure 3), while AUPRC of the second-best method (REVEL) is 0.63, 0.74, 0.73, and 0.35, respectively. PrimateAI, a recent deep representation learning-based method, has a AUPRC of 0.32, 0.68, 0.45, and 0.20, respectively. A comparison using receiver operating characteristic (ROC) curves shows similar patterns (Supplementary Figure 4).

**Figure 3.**
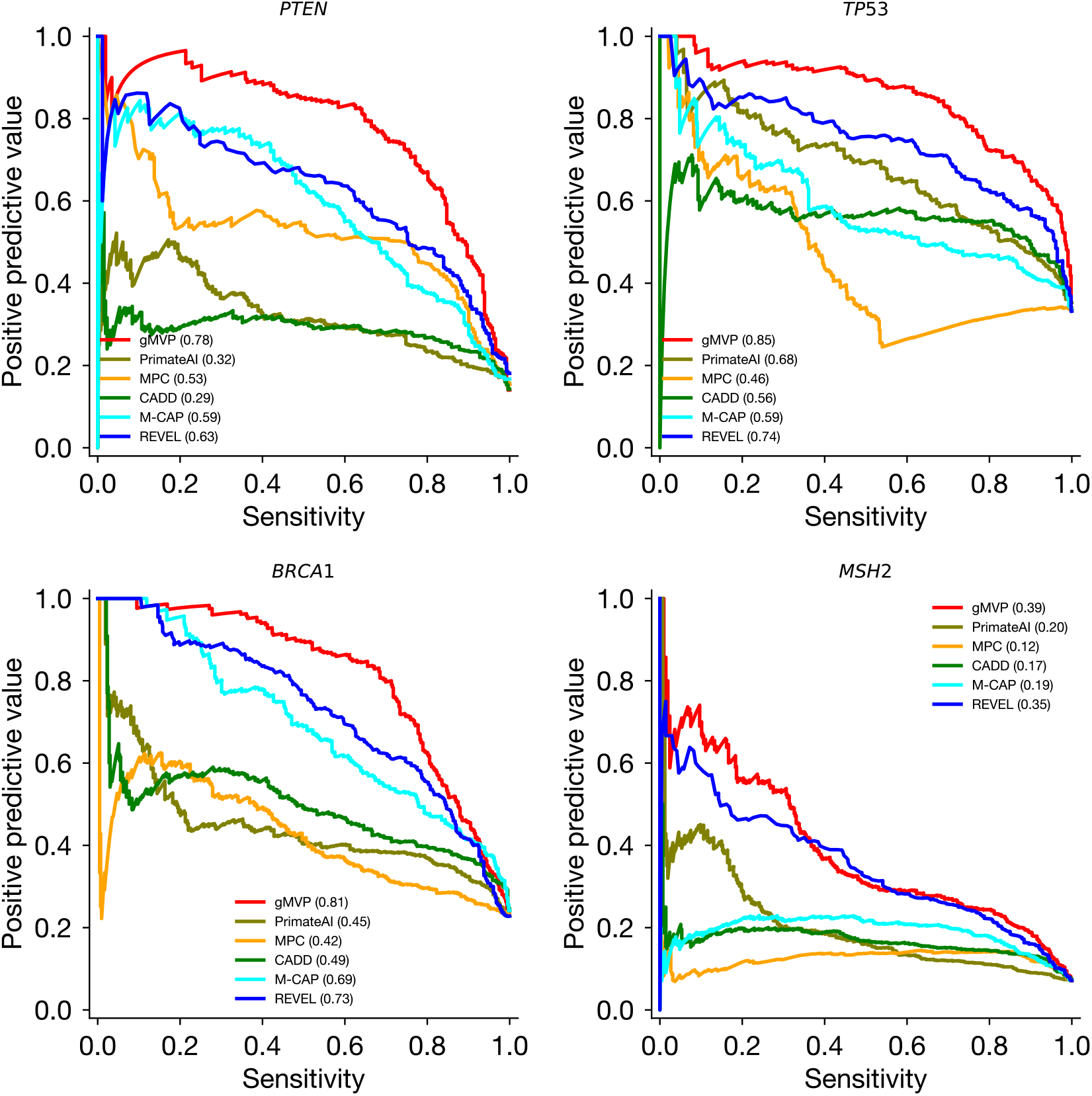
Evaluating gMVP and published methods in identifying damaging variants in known disease genes including *TP53, PTEN, BRCA1*, and *MSH2*. The precision-recall curves of gMVP and published methods are shown for each gene using functional readout data as ground truth.

### Prioritizing rare de novo missense variants in autism spectrum disorder and neural developmental disorders using gMVP

To further evaluate the utility of gMVP in new risk gene discovery, we compared gMVP scores of *de novo* missense variants from cases with developmental disorders and controls. We obtained published *de novo* missense variants (DNMs) from 5924 cases in an autism spectrum disorder (ASD) study^4^, 31058 cases in a NDD study^5^ and DNMs from 2007 controls (unaffected siblings)^4^. Although there is no ground truth because most of these *de novo* variants are not previously implicated with diseases, there is a significant excess of such variants in cases compared to controls^43-45^, indicating that a substantial fraction of variants in cases are pathogenic. We therefore tested whether the predicted scores of variants in cases and controls are significantly different and use significance as a proxy of performance (Figure 4a). gMVP achieves a *p*-value of 3e-9 and 2e-40 for ASD versus controls and NDD versus controls, respectively, while the second-best method PrimateAI achieves a *p*-value of 3e-6 and 2e-38, respectively (Supplementary Fig. 5).

**Figure 4.**
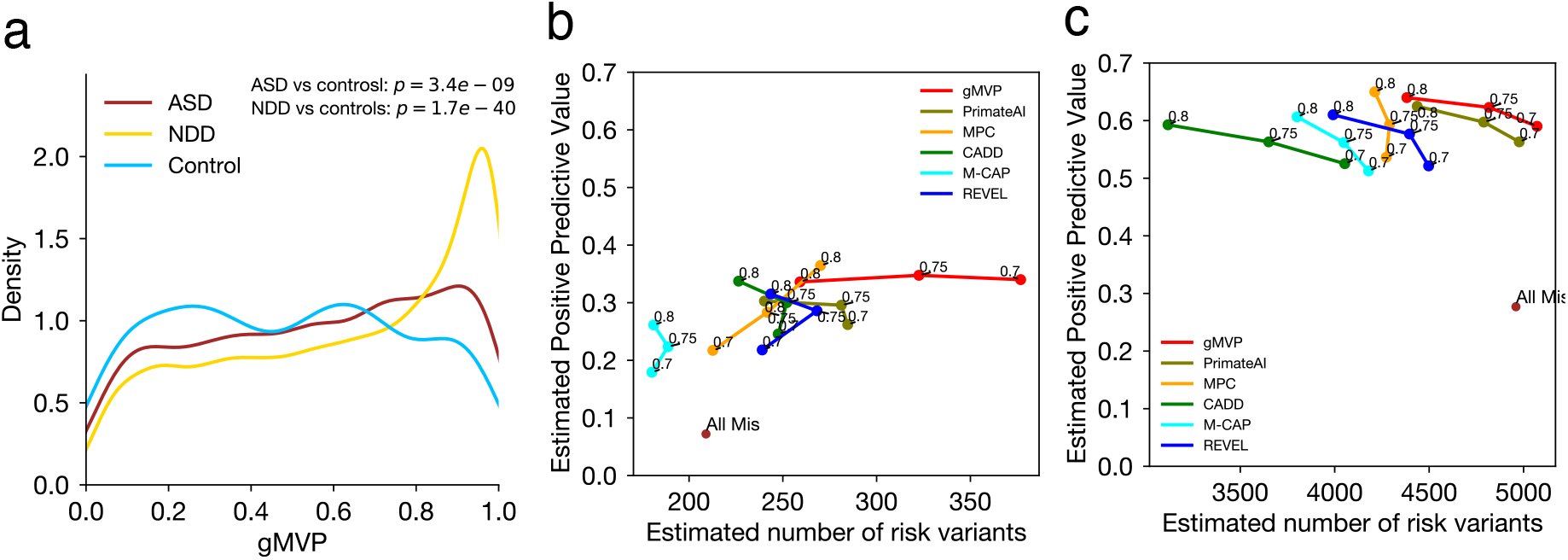
Evaluating gMVP and published methods in distinguishing rare *de novo* missense variants in cases with neurodevelopmental disorders from the ones in controls. **(a)** Distributions of gMVP predicted scores of rare *de novo* missense variants from ASD and NDD cases and controls. We used Mann–Whitney U test to assess the statistical significance of the difference between cases and controls. NDD: neural developmental disorders; ASD: autism spectrum disorder; controls: unaffected siblings from the ASD study. **(b)** Comparison of gMVP and published methods using *de novo* variants from ASD cases and controls by precision-recall-proxy curves. Numbers on each point indicate rank percentile thresholds. The positions of “All Mis” points are estimated from all missense variants without using any prediction method. (**c**) The same comparison using data from NDD cases and controls.

We then calculated the enrichment rate of predicted damaging DNMs by a method with a certain threshold in cases compared to the controls, and then estimated precision and the number of true risk variants (Methods), which is a proxy of recall since the total number of true positives in all cases is a (unknown) constant independent of methods.

The estimated precision and recall values are directly related to power of detecting new risk genes ^5,46^. We compared the performance of gMVP to other methods by estimated precision and recall-proxy (Figure 4b and 4c). The optimal threshold of gMVP rank score in cancer hotspots is 0.75. With 0.75 as the threshold, we observed an enrichment rate of 2.7 in NDD and an enrichment of 1.5 in ASD (Supplementary Table 7 and 8), corresponding to estimated precision-recall of (0.62, 4818) and (0.35, 328), respectively. Additionally, when using a lower threshold 0.7, gMVP can still keep the precision as high as 0.34 and achieved a recall of 377 in ASD. PrimateAI achieved overall second-best estimated precision and recall under different thresholds in both ASD and NDD. MPC with a threshold of 0.8 can reach a high precision at 0.65 and 0.36 in NDD and ASD respectively, but overall it has substantially lower recall than gMVP and PrimateAI.

### Classifying gain-of function and loss-of-function variants using transfer learning

In many genes, the functional impact of missense variants is complex and cannot be simply captured by a binary prediction. Recently, Heyne e*t al*^*47*^ investigated the pathogenetic variants that alter the channel activity of voltage-gated sodium (Navs) and calcium channels (Cavs) and inferred loss-of function (LOF) and gain-of function (GOF) variants based on clinical phenotypes of variant carriers and electrophysiology data. Additionally, the study described a computational model (“funNCion”) to predict LOF and GOF variants using a large number of human-curated features on biochemical properties. Here we sought to classify LOF and GOF variants using gMVP model through transfer learning without additional curated prediction features. Transfer learning allows us to further train a model for a specific purpose using a limited number of training points by only exploring a reasonable subspace of the whole parameter spaces guided by previously trained models.

We obtained 1517 pathogenetic and 2328 neutral variants in 10 voltage-gated sodium and 10 calcium channel genes, in which 518 and 309 variants were inferred as LOF and GOF variants, respectively, from Heyne *et al* ^47^. To benchmark the performance, we used the same training and testing sets (90%/10% breakdown) as funNCion.

We first evaluated the performance of gMVP and previous methods in distinguishing LOF or GOF from neutral variants. gMVP and REVEL both achieved the best AUROC at 0.94 (Figure 5a and Supplementary Table 9). FunNCion^47^ which was trained specifically on the variants of the ion channel genes achieved nearly identical AUROC (0.93). We next sought to improve the performance using transfer learning. Starting from the weights from the original gMVP model, we trained a new model, gMVP-TL1, with both LOF and GOF variants in these genes as positives and neutral variants as negatives (Methods). gMVP-TL1 achieved an AUROC of 0.96, outperforming the original gMVP and published methods. Furthermore, to distinguish LOF and GOF variants, we trained another model, gMVP-TL2, also starting from the weights of the original gMVP model but with different output labels for training (LOF versus GOF) (Methods). The training set includes 465 LOF and 279 GOF variants and the testing set includes 51 LOF and 30 GOF variants. gMVP-TL2 achieved an AUROC of 0.95, substantially better than funNCion (AUROC, 0.84) which trained on the same variants set with manually curated prediction features (Figure 5b and Supplementary Table S10). This demonstrates that the gMVP model aided by transfer learning technique can accurately predict GOF and LOF variants in channel genes with a very limited training dataset.

**Figure 5.**
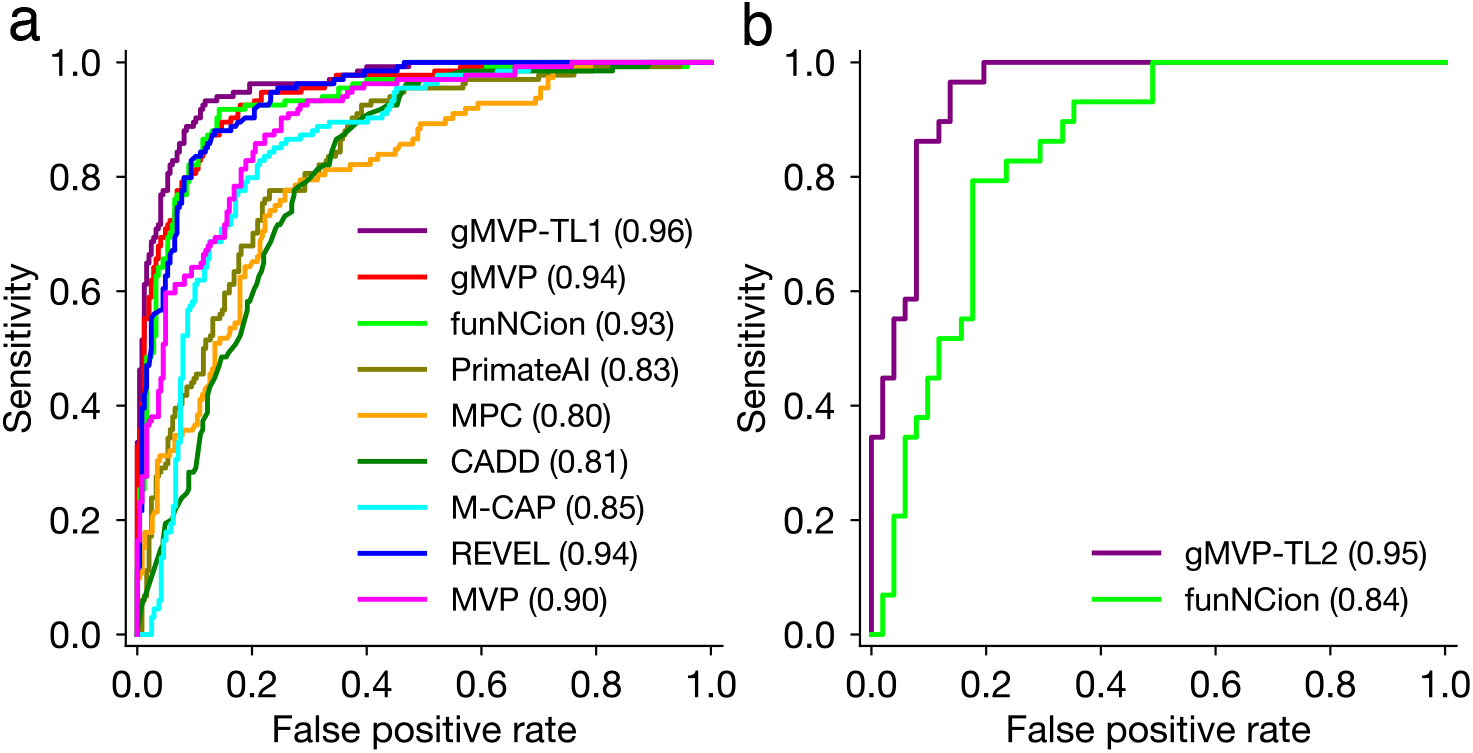
Evaluating gMVP and published methods in classifying pathogenetic and neutral variants and in predicting GOF and LOF variants in ion channel genes. **(a)** Comparison of ROC curves in classifying pathogenetic variants and neutral variants. gMVP-TL1 denotes the model further trained on the pathogenetic and neutral variants in *SCNxA* genes starting from the weights of the original gMVP model. **(b)** Comparison of ROC curves in classifying GOF and LOF variants. gMVP-TL2 denotes the model further trained on GOF and LOF variants starting from the weights of the original gMVP model.

### gMVP prediction captures information on conservation, protein structure, and selection in human

We calculated the correlation between predicted scores of gMVP and other methods on *de novo* variants from ASD and NDD cases and controls (Figure 6a). gMVP has the highest correlation with REVEL (Spearman ρ=0.78), followed by a few other widely used methods such as MPC, CADD, and PrimateAI (ρ>0.6).

**Figure 6.**
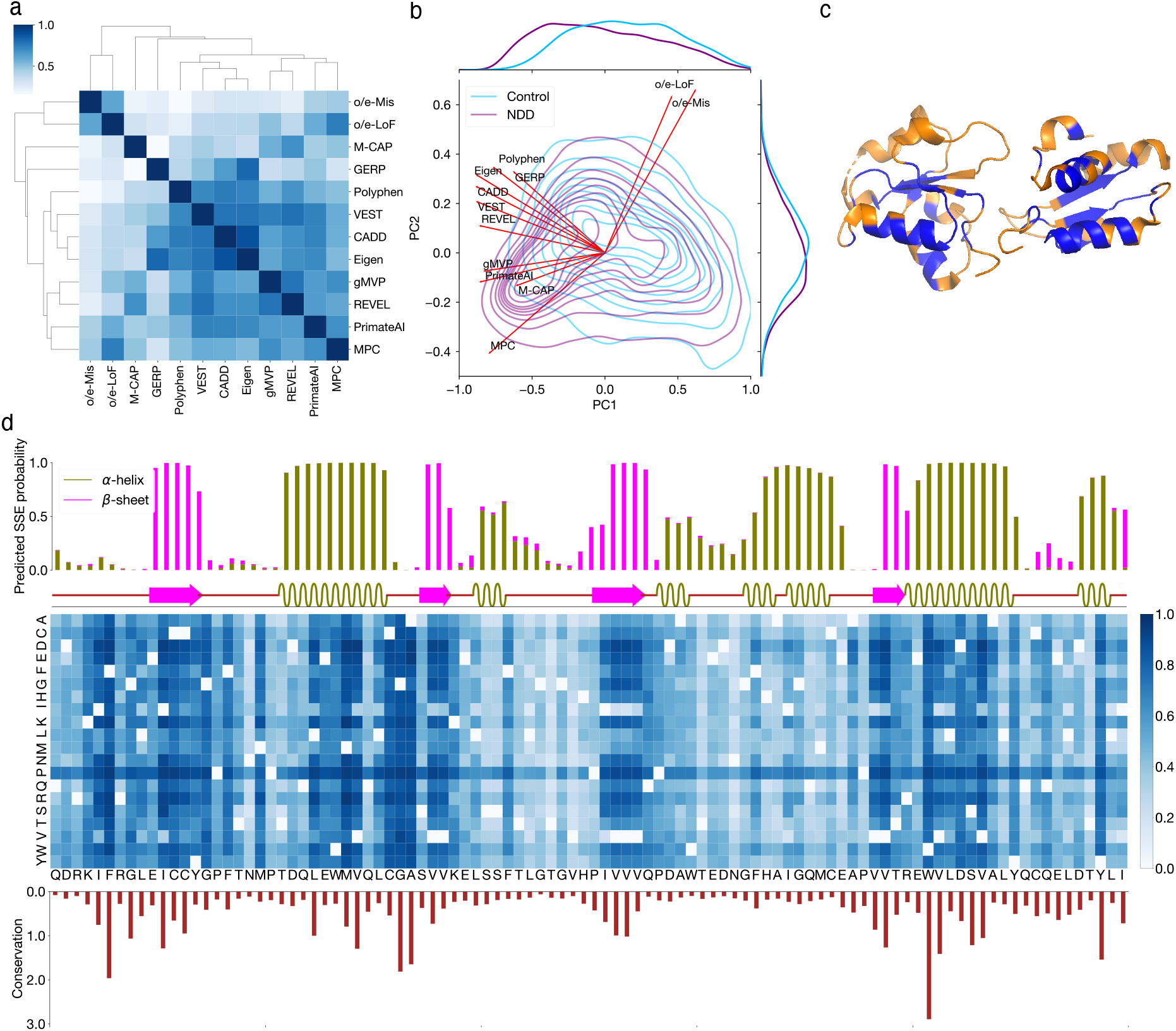
Interpreting gMVP predictions with conservation, protein structure, and genetic coding constraints. (**a**) Spearman correlation between gMVP and other published methods, calculated by scores of the *de novo* variants in ASD, NDD, and controls. (**b**) PCA on *de novo* variants from ASD and NDD cases and controls. Red arrows show the loadings of gMVP and published methods on the first two components; the density contour shows the distribution of PC1/2 scores of the variants in NDD (purple) and controls (light blue). The density curves along the axes show the distribution of PC1 or PC2 scores of the cases and controls. (**c**) The protein tertiary structure of BRCT2 domain of BRCA1. We colored a residue as blue if at least one missense on this position is predicted as damaging (gMVP > 0.75) and orange otherwise. (**d**) gMVP scores of all possible missense variants on *BRCT2* domain of *BRCA1*. The top histogram and the following bar show the predicted and real protein secondary structures, respectively. The middle heatmap shows gMVP scores for all possible missense variants on each protein position. The bottom histogram shows the evolutionary conservation measured with the entropy of the amino acid distribution among homologous sequences.

We then performed principal component analysis (PCA) on the *de novo* variants from cases and controls to investigate the contributing factors that separate the variants in cases and controls (Figure 6b and Supplementary Fig.6). The input of the PCA is a score matrix where rows represent variants and columns represent predicted scores by gMVP and other methods. We included two additional columns with gene-level gnomAD constraint metrics o/e-LoF and o/e-Mis^48^ (observed over expected for LoF and missense) to represent selection effect in human population. The first component (PC1) captures the majority of the variance of the data and best separates the *de novo* variants in cases and the ones in controls. All methods have large loadings on PC1 (Figure 6b). The second component (PC2) is largely driven by the gene-level gnomAD constraint metrics (Figure 6b). The joint distribution of PC1/2 scores of DNMs from controls has a single mode at the center. The joint distributions of scores of DNMs from cases have two modes (Figure 6b and Supplementary Fig. 6b), representing mixtures of likely pathogenic variants and random DNMs. Notably, gnomAD metrics have near orthogonal loadings on PC1/2 with GERP which is purely based on cross-species conservation, suggesting that selection effect in human provides complementary information to evolutionary conservation about genetic effect of missense variants. All methods (PolyPhen, eigen, CADD, VEST, and REVEL) that do not use human or primate population genome data have loadings close to GERP on PC1/2. MPC and M-CAP, which use sub-genic or gene-level mutation intolerance metrics similar to gnomAD metrics, have closest loadings as gnomAD metrics on PC1/2. gMVP and PrimateAI have similar loadings that are in the middle of GERP and gnomAD metrics.

We inspected the *BRCT2* domain of *BRCA1* to show how the gMVP model captures context-dependent functional impact. We observed that most damaging variants predicted by gMVP (>0.75) are located in the core region of *BRCT2* domain (Figure 6c). Additionally, gMVP scores are highly correlated with evolutionary conservation (Figure 6d and Supplementary Fig. 7a, ρ=0.57). Variants in the *β*-sheets are significantly more damaging than the ones in *α*-helix regions, and the ones in *α*-helix regions are more damaging than the ones in coil regions (Figure 6d and Supplementary Fig. 7b), consistent with previous discoveries^21,49,50^. Finally, amino acids mutated to *Proline* (P) in helix regions are predicted to be highly damaging, even in positions not well conserved (Figure 6d). This is consistent with the fact that *Proline* rarely occurs in the middle of an alpha-helix^51^.

## Discussion

We developed gMVP, a new method based on graph attention neural networks, to predict functionally damaging missense variants. gMVP uses attention neural networks to learn representations of protein sequence and structure context through supervised learning trained with large number of curated pathogenic variants. The graph structure allows coevolution-guided pooling of predictive information of distal amino acid positions that are functionally correlated or potentially close in 3-dimensional space. We demonstrated the utility of the gMVP in clinical genetic testing and new risk gene discovery studies. Specifically, we showed that gMVP achieves better accuracy in identification of damaging variants in known risk genes based on functional readout data from deep mutational scan studies. Additionally, gMVP achieved better performance in prioritizing *de novo* missense variants in cases with autism or NDD, suggesting that it can be used to pre-select damaging variants or weight variants to improve statistical power of new gene discovery. Finally, we showed that with transfer learning technique, gMVP model can accurately classify GOF and LOF variants in ion channels even with a limited training set without additional prediction features.

gMVP learns a representation of protein context from training data, while previous ensemble methods such as REVEL, M-CAP, MetaSVM, and CADD used scores from other predictors or other human-engineered features as inputs. With recent progress of machine learning in protein structure prediction ^52-55^, neural network representations could capture latent structure beyond common linear representations of understanding of the biophysical and biochemical properties. We showed that representation learning allows gMVP to capture the context-dependent impact of amino acid substitutions on protein function. PrimateAI is a recently published method that also uses deep representation learning. gMVP achieved better performance than PrimateAI in identification of damaging variants in known disease risk genes in comparisons using functional readout data and in prioritizing rare *de novo* variants from ASD and DDD studies. While both models used evolutionary conservation and protein structural properties as features, the two methods have entirely different model architecture and training data. gMVP uses a graph attention neural network to pool information from both distal and local positions with coevolution strength, while PrimateAI uses a convolutional neural network to extract local patterns from protein context. For training data, gMVP used expert-curated variants and random variants in population as training positives and negatives, respectively. In contrast, PrimateAI used common variants in primates as negatives and unobserved variants in population as positives. Based on functional readout data of the four well-known risk genes, only 15-25% of random variants have discernable impact on protein function. Therefore, the positives used in PrimateAI training may contain a large fraction of false positives. PrimateAI’s training strategy does have advantages. It avoids human interpretation bias and errors in curated databases of pathogenic variants, the positives used in gMVP training. It also can cover almost all human protein-coding genes, whereas curated databases such as ClinVar only cover hundreds of genes. Additionally, common variants in primates are likely all true negatives, whereas random observed rare variants in human population could have a non-negligible fraction of damaging variants. Making a new model that can utilize all these datasets in training could further improve the prediction performance.

Several previous studies have shown that the functional impact of missense variants is correlated among 3-dimensional neighbors^21,22,56^. Pooling information from 3-dimensional neighbors could therefore improve predictions of functional impact. However, directly considering 3 dimensional distances is limited by the fact that most human proteins have no solved tertiary structures with considerable coverage. gMVP addresses this issue by taking a large segment of the protein context that include both local and distal positions that are potential neighbors in folded proteins, and then uses coevolution strength to effectively pool information from potential 3D neighbors. Used as edge features in a graph attention model, coevolution strength allows more precise pooling of information from distal residues than the convolutional layer without prior structure. Coevolution strength has been used in *ab initio* protein structure prediction extensively^30,55,57^. The extraordinary performance of AlphaFold2^53^ in CASP14 shows that it contains critical information about physical residue-residue distances for accurate structure prediction to many more proteins. More recently, the language model Transformer^33^ has been applied on protein sequences and multi-sequence alignments (MSAs) to improve the performance of coevolution strength estimation and protein residue-residue contacts prediction^58-60^. gMVP could be further improved by integrating components of Transformer in the model.

With transfer learning, the trained gMVP model can be further optimized for more specific tasks in genetic study. The idea is to transfer the general knowledge learned from large training data sets to a new related and more specific task with only limited training data. The trained model can set the initial values of the weights in the model to be updated by further training to explore only a subspace of the whole parameter space. We have shown its feasibility in classifying GOF and LOF variants in the ion channel genes using a limited number of training data points without additional prediction features. We expect that with transfer learning, gMVP can potentially improve variant interpretation by training on gene family-specific models^61^ and to identify disease-specific damaging variants^62^.

Functional readout data from deep mutational scan provides strong evidence of classifying variants as damaging or neutral^25-28,63,64^. However, these *in vitro* functional readout assay usually reveals only one aspect of a protein’s function in a limited number of cell types, therefore, they are often not completely correlated with the functional impact of the variants *in vivo*. We expect that more comprehensive deep mutational scan assays will become available and facilitate substantial improvement in the training and evaluation of computational methods.

Finally, we showed that while evolutionary conservation remains one of the most informative sources for computational methods, selection in human population can provide complementary information for prediction. Selection coefficient is correlated with allele frequency, especially for variants under strong negative selection ^46,65-67^. Larger population genome data sets can further improve estimation of allele frequency of rare variants. We anticipate large ^68^ and diverse ^69^ population genome data released in the future will improve estimation of selection effect in human and in turn improve gMVP.

## Methods

### Training data sets

For positive training set, we collected 22,607 variants from ClinVar database^37^ under the Pathogenic and Likely-Pathogenic categories with review status of at least one star, 48,125 variants from Human Gene Mutation Database Pro version 2013 (HGMD) database^36^ under the disease mutation (DM) category, and 20,481 variants from UniProt labeled as Disease-Causing. For negative training sets, we collected 41,185 variants from ClinVar under the Benign and Likely-Benign categories, 33,387 variants from SwissVar^38^ labeled as Polymorphism. After excluding 3,751 variants with conflicting interpretations by the three databases, we have 63,304 and 66,102 unique positives and negatives. We next excluded 36,499 common variants (653 positives and 35,846 negatives) with allele frequency > 1e-3 in gnomAD (all populations) ^70^ and 3,080 overlapping variants (2,680 positives and 400 negatives) with testing datasets from the training dataset, resulting in a dataset of 59,701 positives and 29,856 negatives. To balance the positive and negative training samples, we randomly selected 29,845 rare missense variants from DiscovEHR database^42^ that are not already covered by previously selected training data as additional negative training points. In the end, we have 59,701 and 59,701 unique positive and negative training variants (Supplementary Table 1), which cover 3,463 and 14,222 genes, respectively.

### Testing data sets

1. Cancer somatic mutation hotspots: we obtained 878 missense variants located in somatic missense mutations hotspots in 209 cancer driver genes from a recent study^24^ as positives, and randomly selected 2 times more rare missense variants (N=1756) from the population sequencing data DiscovEHR^42^.
2. Functional readout data from deep mutational scan experiments: we compiled variants in *BRCA1*^26^, *PTEN*^27^, *TP53*^28^, and *MSH2*^*25*^. We only include the single nucleotide variants (SNVs) for comparison as most published methods don’t provide scores for the non-SNVs. There are 432 positives and 1,476 negatives in *BRCA1*, 258 positives and 1601 negatives in *PTEN*, and 540 positives and 1,108 negatives in *TP53*, and 414 positives and 5439 negatives in *MSH2*.
3. *De novo* variants: to evaluate utility in new risk gene discovery, we used published rare germline *de novo* missense variants (DNVs) from 5,924 cases and 2,007 controls in an autism spectrum disorder (ASD) study^4^ and 31,058 cases in a neural developmental study^5^.

To fairly compare our methods with published methods, we excluded the overlapping variants with testing datasets from the training datasets. We further excluded all variants in *PTEN, TP53, BRCA1*, and *MSH2* in training to avoid inflation in performance evaluation.

### The Graph Attention Neural Network model

gMVP uses a graph to represent a variant and its protein context. We first defined the 128 amino acids flanking the reference amino acid as protein context. We next built a star-like graph with the reference amino acid as the center node and the flanking amino acids as context nodes, and with edges between the center node and each context node (Figure 1 and Supplementary Fig. 1).

Let ***x, n***_*i*_, and ***f***_***i***_ denote input feature vectors for the center node, each context node, and each edge, respectively. We first used three 1-depth dense layers to encode ***x, n***_*i*_, and ***f***_***i***_ to latent representation vectors ***h, t***_***i***_, and ***e***_***i***_, respectively. We used RELU^71^ as the activation function and 512 neurons for each dense layer.

We then used a multi-head layer adapted from the attention layer in the Transformer model^33^ to pool information from context nodes and finally to learn a context vector ***c***. Specifically, for the *k*th head, we first calculated the value vectors for each context node by 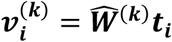 We next calculated attention scores for each context node through 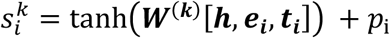 where *tanh* denotes a hyperbolic tangent activation function, and *p*_i_ is a position bias which is a simplified positional encoding^72^. *We note here p*_*i*_ *allows the model to capture local protein sequence context*. Attention weights are calculated by applying a *softmax* operation on the attention scores, 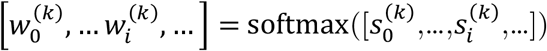.

The context vector ***c***^(*k*)^ for the *k*th head is calculated as 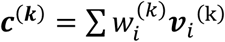. The final context vector is obtained by a linear projection on the concatenation vector of the context vectors from each head,

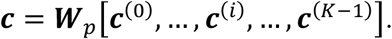

Here *K* denotes the number of heads and we used 4 heads in our model. And we note that in the model, 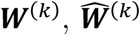, and ***W***_*p*_ are weight matrices to be trained.

We next used a gated recurrent unit (GRU) layer^35^ to leverage the context vector ***c*** and the latent vector ***h*** of the given variant where the relative importance of the whole context can be determined. We used 512 neurons and a hyperbolic tangent activation function for the GRU layer. We finally used a linear projection layer and a sigmoid layer to perform classification.

### Input features

The center node, which represents the variant, has the following features: reference and alternate amino acids, evolutionary conservation, and predicted local structural properties. The context nodes have the following features: reference amino acids, evolutionary conservation, predicted local structural properties, and observed and expected missense alleles in gnomAD ^48^. The feature of edges is coevolution strength between the position of variant and other positions, estimated from multiple sequence alignments of homologous sequences.

#### *Reference and alternate amino acids* (40 values)

we used one-hot encoding with a dimension of 20 to represent reference and alternate amino acids.

#### *Protein primary sequence* (20 values)

We also used one-hot encoding to represent each amino acid in the protein primary sequence.

#### *Evolutionary conservation* (42 values)

we estimated the evolutional conservation from two sources: (1) we searched the homologous of the protein of interest against SwissProt database^73^ with 3 iterations of search and then built the multiple sequence alignments (MSAs) with HHblits suite^74^. (2) we downloaded the MSAs of 200 species from Ensembl website for each human protein sequence^75^. We then calculated the frequencies of 20 amino acids and the gap for each position for the two MSAs separately and concatenated the two frequency vectors.

#### *Predicted protein structural properties* (5 values)

we predicted the protein secondary structures (3 values), solvent accessibility (1 value), and the probability of a residue participating in interactions with other proteins (1 value) using NetsurfP^76^.

#### *Observed number of missense alleles in gnomAD and expected number* (2 values)

to capture selection effect in human, we obtained the observed number of rare missense variants in gnomAD^48^ and the expected number of rare missense variants estimated using a background mutation model^48^.

#### *Coevolution strength* (442 values)

We extract pairwise statistics from the MSA as coevolution strength. It is estimated based on the covariance matrix constructed from the input MSA. First, we compute 1-site and 2-site frequency counts 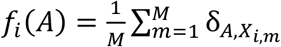 and 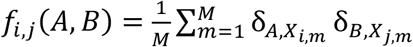, where A and *B* denote amino acid identities (20 + gap), δ is the Kronecker delta, *i* and *j* are position indexes on the aligned protein sequence, *m* is the sequence index of the MSA with a total of *M* aligned sequences, and *X*_*i,m*_ indicates the amino acid identity of position *i* on sequence *m*. We then calculate the sample covariance (21×21) matrix 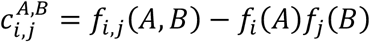 and flatted it into a vector with 441 elements. We also convert the covariance matrix to a single value by computing its Frobenius norm 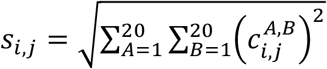 and then concatenate the norm and the flattened vector as the edge features.

We built these features only for canonical transcripts defined by Ensembl ^77^ Version92. We annotated the variants using VEP^78^.

### Training algorithm

We used cross-entropy loss as the training loss. We used the Adam algorithm^39^ to update the model parameters with an initial learning rate of 1e-3 and decayed the learning rate with a polynomial decay schedule^79^. We randomly selected 10% of training samples as validation set and early stopping was applied with validation loss as watching metric. We trained 5 models by repeating the above training process five times and for testing we averaged the outputs of the five models as prediction scores. The model and training algorithm were implemented using TensorFlow^40^.

## Classifying GOF and LOF variants using gMVP model and transfer learning

To investigate the potential for transfer learning, we further trained gMVP to classify GOF and LOF variants in *ion* channel genes with additional training data but without new features. We collected 1517 pathogenetic and 2328 neutral variants in *SCNxA* genes which encode Voltage-gated sodium (Navs) and calcium channels (Cavs) protein, in which 518 and 309 variants are inferred as LOF and GOF variants, respectively, from a recent study^47^.

We first trained a model, gMVP-TL1, to classify pathogenetic and neutral variants in *SCNxA* genes. We used the same data set as funNCion^47^, including 3466 variants for training and 379 variants for testing. We randomly selected 10% variants from training set as validation set. We used the same model architecture with gMVP and the weights of gMVP model previous trained using all genes as the initial values of new model. In the new model training, we used Adam algorithm to update parameters with an initial learning rate of 1e-3, and used the validation loss as stopping criteria. We trained 5 gMVP-TL1 models, starting from each of the 5 trained gMVP models and for testing we averaged the outputs of these models as prediction scores.

We next trained another model gMVP-TL2 to classify GOF versus LOF variants in *SCNxA* genes. We used 744 variants as training set and 81 variants as testing set, which are same sets used by funNCion^47^. Like gMVP-TL1, gMVP-TL2 were also trained staring from the weights of gMVP model previous trained using all genes. We used the same hyperparameter setting with gMVP-TL1 in training.

### Normalization of scores using rank percentile

For each method, we first sorted predicted scores of all possible rare missense variants across all protein-coding genes, and then converted the scores into rank percentiles. The higher rank percentile indicates more damaging, e.g., a rank score of 0.9 indicates the missense variant is more likely to be damaging than 90% of all possible missense variants.

### Precision-recall-proxy curves

Since there is no ground truth data to benchmark our performance on *de novo* variants, we estimate precision and recall at various thresholds based on the enrichment of predicted damaging variants in cases compared to controls.

Let *S*_1_ be the rate of synonymous variants in cases, and *S*_0_ be the rate of synonymous variants in controls. Then the synonymous rate ratio α is defined as

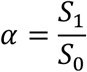

Denote the total number of variants in cases as *N*_1_, the number of variants in controls as *N*_0_, the number of variants predicted as pathogenic in cases as *M*_1_, and the number of variants predicted as pathogenic in controls as *M*_0_. We assume that for there to be no batch effect, the rate of synonymous variants should be the same in the cases and controls. So, we estimate the enrichment of predicted pathogenic variants in cases compared to controls by:

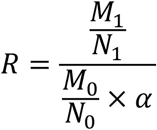

Then, the true number of pathogenic de novo variants 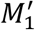 is estimated by

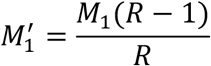

And the estimated precision is

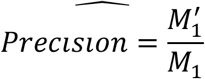

## Data availability

1. Precomputed gMVP scores for all possible missense variants in canonical transcripts on human hg38 can be downloaded from: https://www.dropbox.com/s/nce1jhg3i7jw1hx/gMVP.2021-02-28.csv.gz?dl=0.
2. The training data of the main model were downloaded from: http://www.discovehrshare.com/downloads (DiscovEHR), http://www.hgmd.cf.ac.uk/ac/index.php (HGMD), https://www.uniprot.org/docs/humpvar (UniProt), and https://ftp.ncbi.nlm.nih.gov/pub/clinvar/vcf_GRCh37/ (ClinVar).
3. Other data sets supporting the findings of this study are available in the manuscript and supplementary information files.

## Code availability

The codes for the model design and training and testing procedure are available on GitHub: https://github.com/ShenLab/gMVP/

## Acknowledgements

This work was supported by NIH grants R01GM120609, R03HL147197, and U01HG008680, and Columbia University Precision Medicine Joint Pilot Grants Program. We thank Dr. Xiao Fan, Yige Zhao, Guojie Zhong, Dr. Mohammed AlQuraishi, and Dr. David Knowles for helpful discussions.

## Competing interests

The authors declare no competing interests.

## Notes

### Competing Interest Statement

The authors have declared no competing interest.

### Summary of Updates

Expanded discussions and supplementary materials

