## Supplementary notes for "Predicting functional effect of missense variants using graph attention neural networks"

**Zhang *et al.***

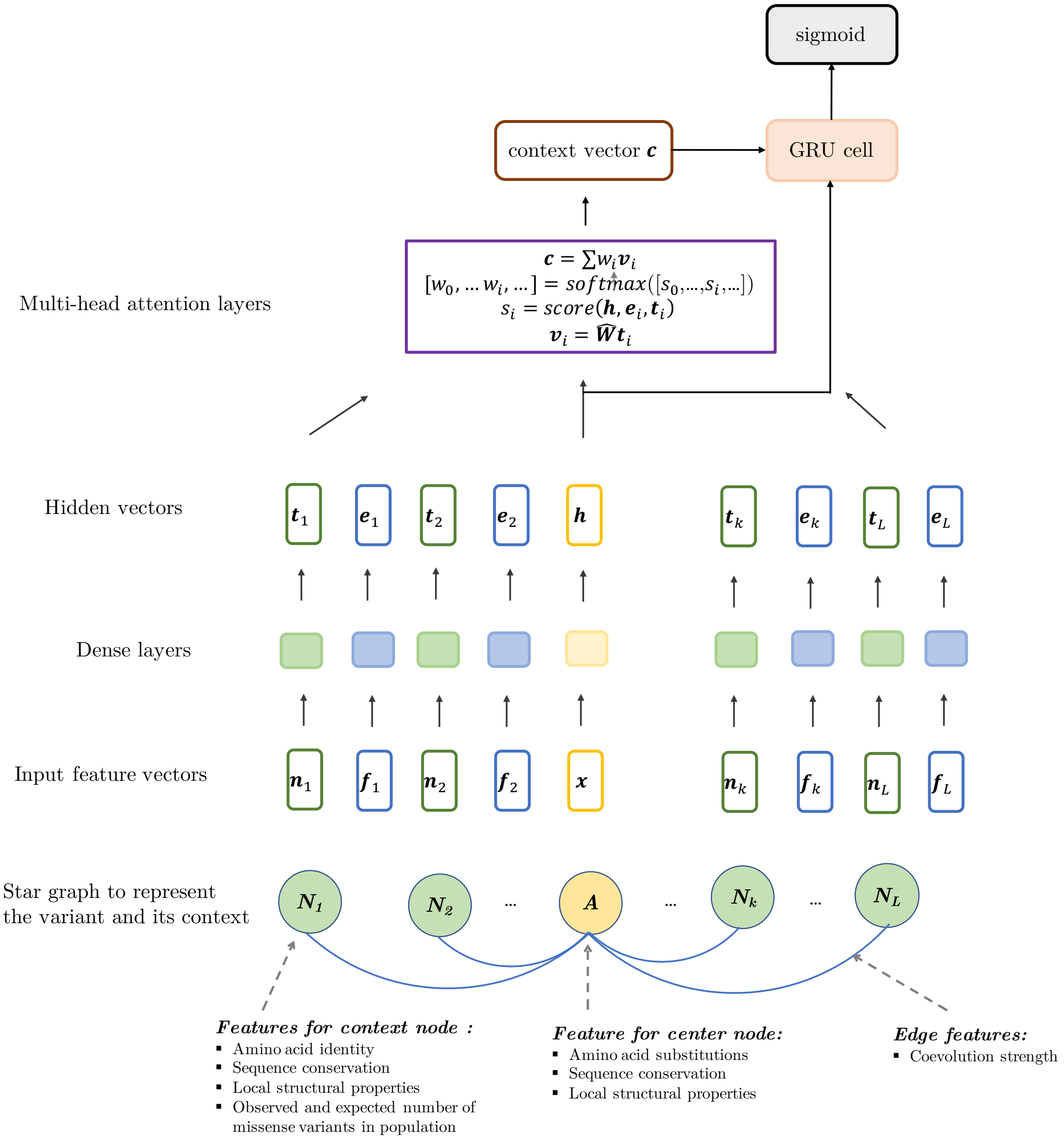

**Supplementary Figure 1. The model architecture of gMVP.** We first build a graph to represent a missense variant and its protein context defined as 128 amino acids flanking the amino acid of interest. The amino acid of interest is the center node (colored as orange) and the flanking amino acids are the context nodes (colored as light green). All context nodes are connected with the center node but not with each other. We use coevolution strength as edge features and used conservation and structural properties as features for both center node and context nodes. We additionally include amino acid substitution as features for center node and primary sequence and the expected and observed number of rare missense variants in the general population for context nodes. The input feature vectors for edges, center node, and context nodes are denoted as $\boldsymbol{f_k}$, $\boldsymbol{x}$, and $\boldsymbol{n_k}$ We apply three 1-depth dense layers to encode the input feature vectors $\boldsymbol{f_k}$, $\boldsymbol{x}$, and $\boldsymbol{n_k}$ to latent vectors $\boldsymbol{e_k}$, $\boldsymbol{h}$, and $\boldsymbol{t_k}$, respectively. We next use a multi-head attention layer to learn a context vector $\boldsymbol{c}$. We then use a gated recurrent neural layer to leverage the context vector $\boldsymbol{c}$ and the latent vector of the variant node $\boldsymbol{h}$. We finally used a sigmoid layer to perform classification.

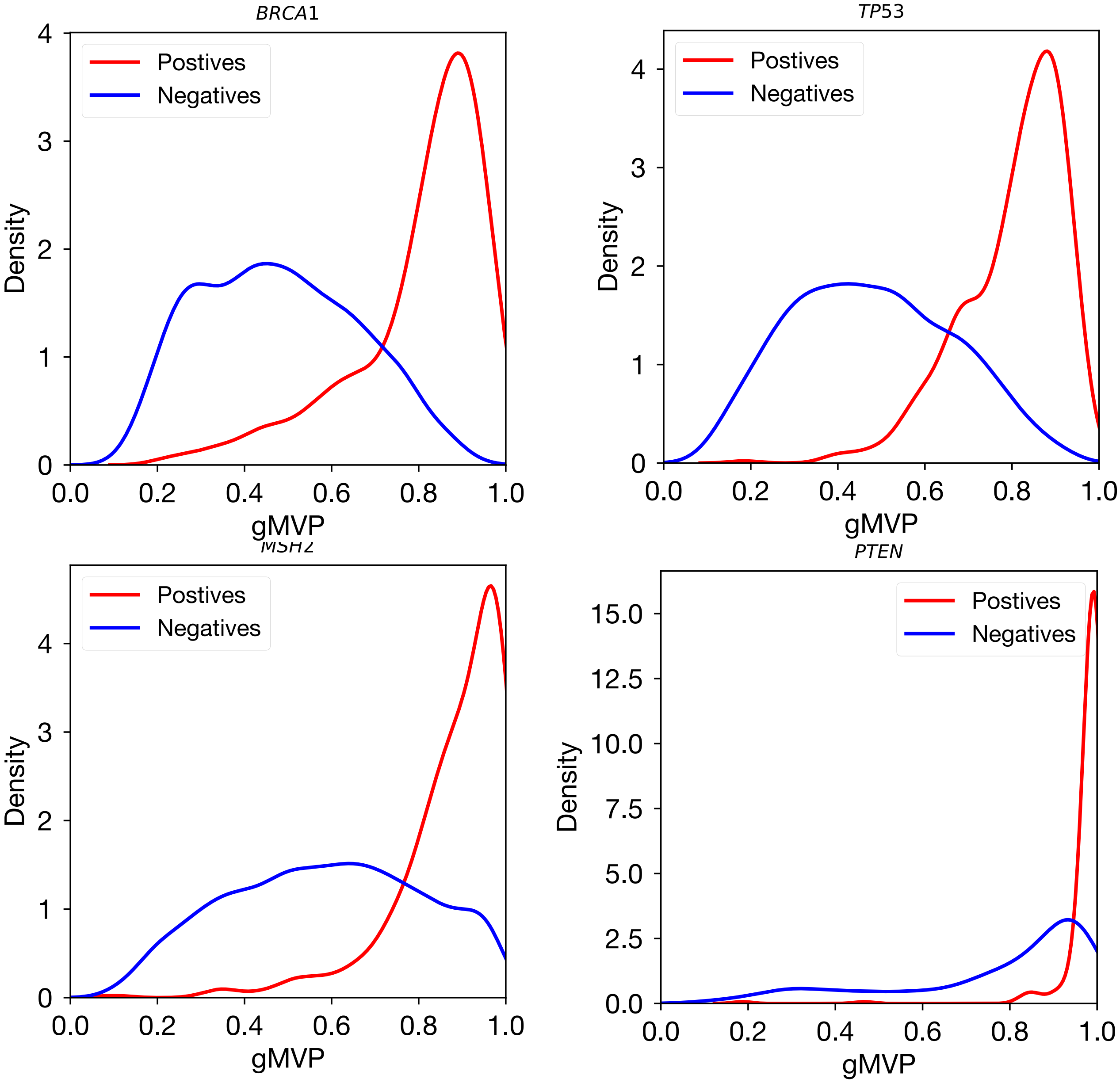

**Supplementary Figure 2. The distributions of gMVP scores** of the damaging *(labeled positive)* and neutral variants *(labeled negatives)* on known disease genes, including *TP53*, *PTEN*, *BRCA1*, and *MSH2*. The labels were determined by functional readout data of deep mutational scan experiments.

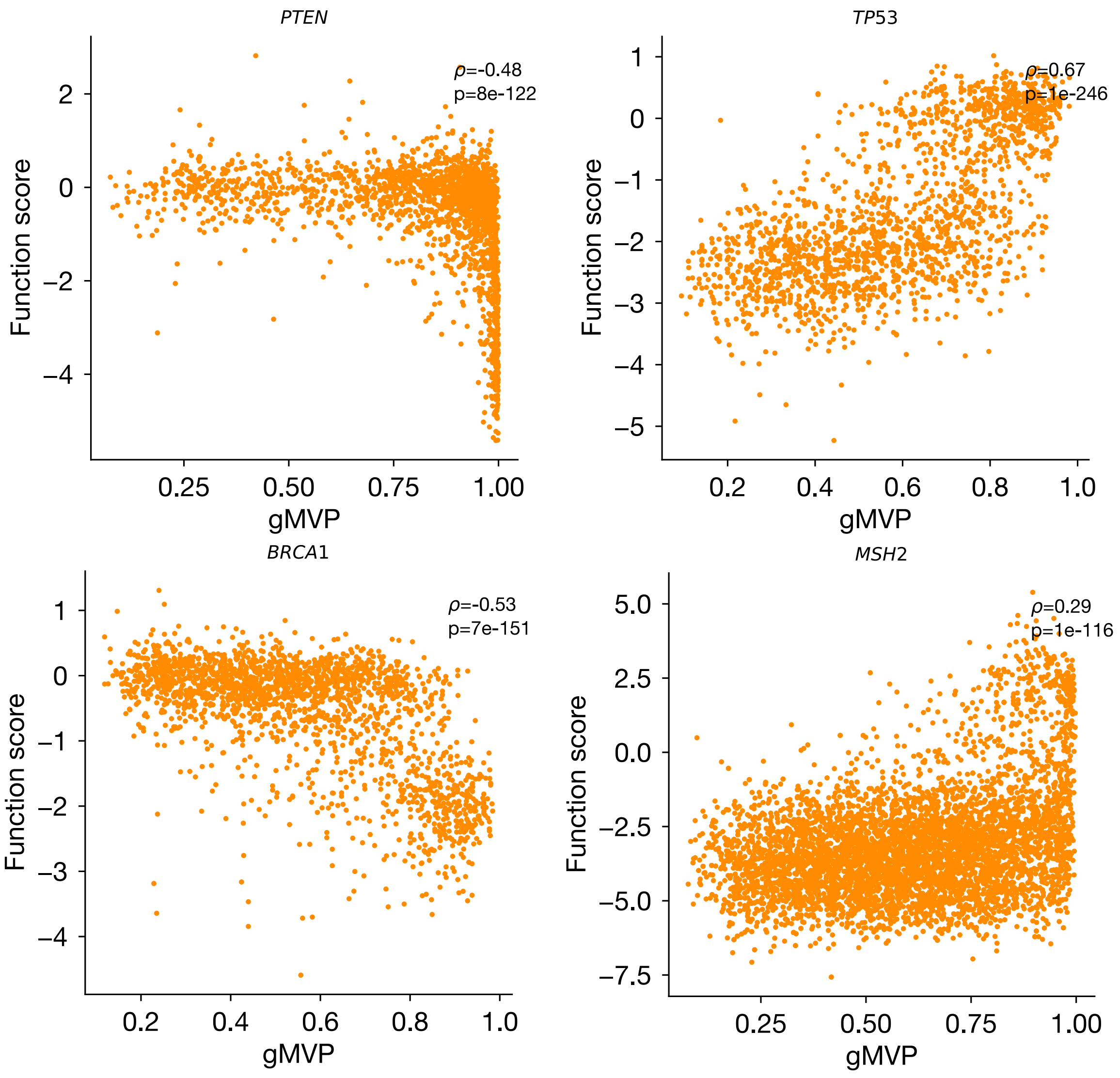

**Supplementary Figure 3.** gMVP scores are correlated with functional readout data from deep mutational scan experiments of known disease genes, including *PTEN*, *TP53*, *BRCA1*, and *MSH2*.

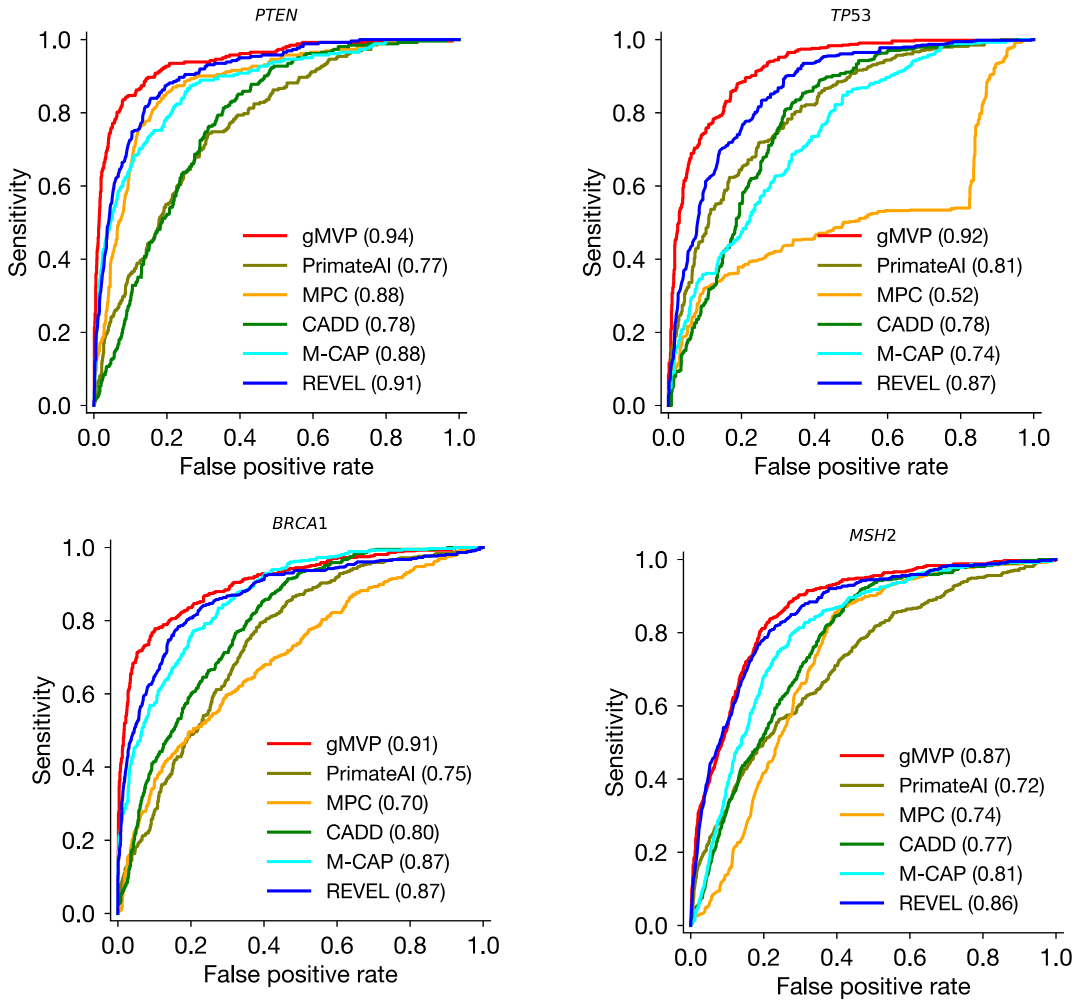

**Supplementary Figure 4. Evaluating gMVP and published methods** in identifying damaging variants on known disease genes, including TP53, PTEN, BRCA1, and MSH2. The receiver operating characteristic curves (ROC) of gMVP and published methods are shown for each gene using functional readout data as ground truth. There are 432 positives and 1476 negatives in BRCA1, 258 positives and 1601 negatives in PTEN, and 540 positives and 1108 negatives in TP53, and 414 positives and 5439 negatives in MSH2.

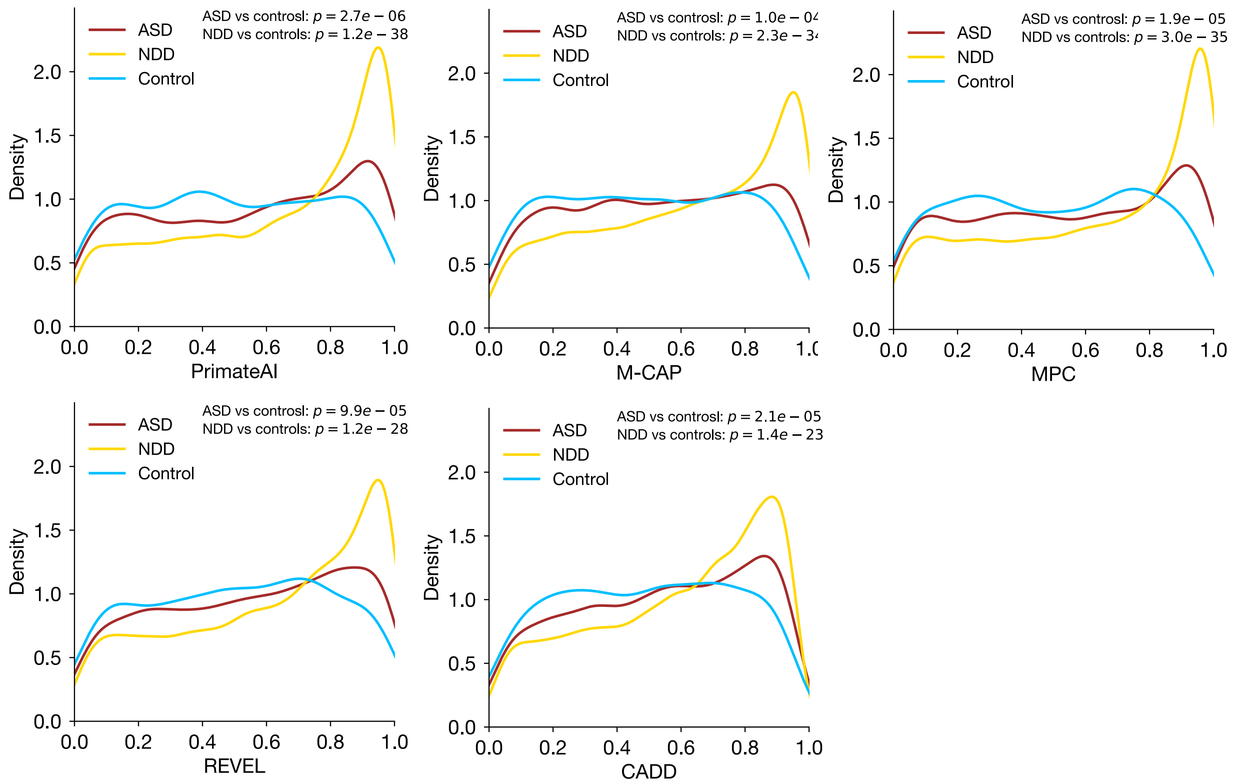

**Supplementary Figure 5. Distributions of predicted scores of published methods of rare *de novo* missense variants from ASD and NDD cases and controls**. We used Mann–Whitney U test to assess the statistical significance of the difference between cases and controls. NDD: neural developmental disorders; ASD: autism spectrum disorder; controls: unaffected siblings from the ASD study. Number of *de novo* missense variants compared: ASD: 2,913; NDD: 17,964; controls: 927. Number of trios in the data: ASD: 5,924; NDD: 31,058; controls: 31058.

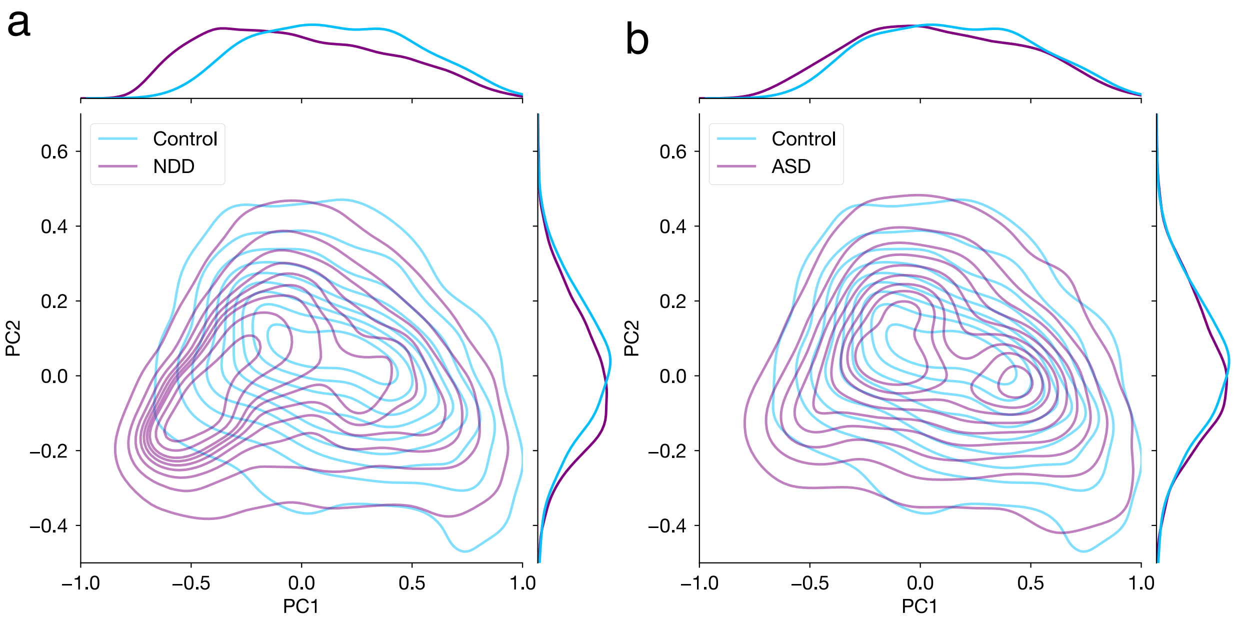

**Supplementary Figure 6. Principal component analysis of *de novo* missense variants in ASD and controls.** We performed principal component analysis (PCA) on the *de novo* variants from cases and controls. The input of the PCA is a score matrix where each row represents a variant and each column represents the predicted score of gMVP or other methods. The density contours show the distribution of PC1/2 scores of the variants in cases and controls. The density curves along the axes show the distribution of PC1 or PC2 scores of cases and controls. **(a)** PC1 versus PC2 of *de novo* variants from NDD cases and controls. **(b)** PC1 versus PC2 of *de novo* variants from ASD cases and controls.

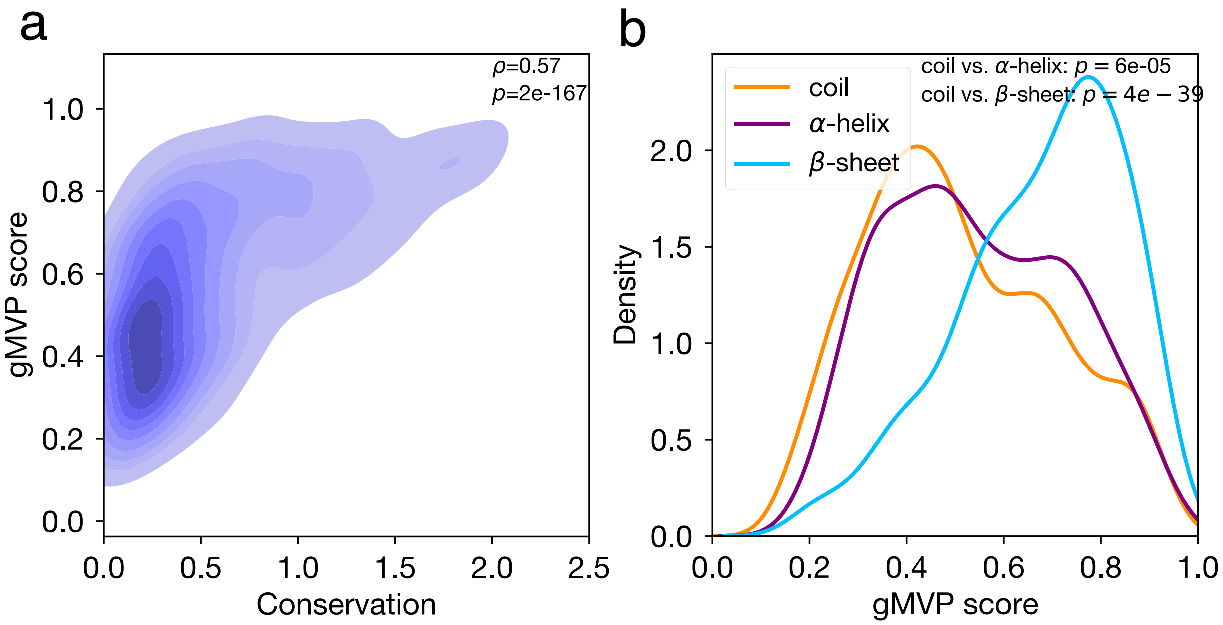

**Supplementary Figure 7. gMVP scores correlate with evolutionary conservation and protein secondary structure**. We show gMVP scores of all possible missense variants in *BRCT2* domain of *BRCA1*. We measured the evolutionary conservation for each protein position with the entropy of the amino acid distribution among homologous sequences. We obtained the secondary structures using the solved protein structure of *BRCT2* domain. (**a**) gMVP scores versus evolutionary conservation. (b) Distributions of gMVP scores of variants located on the coils, $\alpha$-helices, and $\beta$-sheets, respectively. We used Mann–Whitney U test to assess the statistical significance of the difference between the gMVP score of variants on the coils and on the $\beta$-sheet and $\alpha$-helix regions.

### Description of supplementary tables

**S1**. Summary statistics of training data sets.
**S2.** Somatic mutations in cancer hotspots and random variants from DiscovEHR with annotations.
**S3-S6.** Variants with functional readout data from deep mutational scan experiments of *BRCA1*, *TP53*, *MSH2*, and *PTEN*.
**S7.** NDD *de novo* variants enrichment by various methods at rank percentile thresholds
**S8.** ASD *de novo* variants enrichment by various methods at rank percentile thresholds
**S9.** Pathogenetic and neutral variants in ion channel genes and the annotations.
**S10.** GOF and LOF variants in ion channel genes and the annotations.
